# eSkip2 prioritizes exon-skipping antisense oligonucleotide target regions across exon–intron contexts

**DOI:** 10.64898/2026.05.05.722571

**Authors:** Shuntaro Chiba, Katsuhiko Kunitake, Satomi Shirakaki, Umme Sabrina Haque, Harry Wilton-Clark, Md Nur Ahad Shah, Jamie Leckie, Kosuke Matsui, Fumie Uno-Ono, Toshifumi Yokota, Yoshitsugu Aoki, Yasushi Okuno

## Abstract

Exon-skipping antisense oligonucleotides (ASOs) can restore productive transcripts, but identifying effective binding regions remains difficult because splicing regulation extends across exons, introns and splice junctions. Here we develop eSkip2, a genome-informed framework that ranks target regions within a unified exon–intron sequence context. eSkip2 combines a genome-pretrained sequence model with ASO-induced exon-skipping data and single-nucleotide-variant splicing perturbations, followed by target-locus adaptation that requires no experimental ASO labels from the locus being designed. Across benchmarks comprising canonical exons and pseudoexons, multiple cell types and chemistries, and exonic, intronic and exon–intron-spanning targets, eSkip2 prioritized active regions and showed a higher median AUROC than applicable exon-restricted models. Prospective application to the combinatorial design of dual-targeting ASOs for DMD exon 46 enriched active candidates near the top of the ranking: the two most active new ASOs ranked within the top three and produced dose-dependent dystrophin restoration in patient-derived cells. These results support eSkip2 as a practical first-pass strategy for reducing experimental search space in exon-skipping ASO discovery.

## Introduction

Antisense oligonucleotide (ASO)-mediated exon skipping can restore productive transcripts by masking regulatory or splice-junction sequences in pre-mRNA, thereby excluding a selected exon or pathogenic pseudoexon from the mature transcript^1–5^. The strategy is clinically established in Duchenne muscular dystrophy (DMD) and is increasingly relevant to Mendelian disorders caused by deep-intronic variants^4,5^. Frameworks for individualized splice-switching therapy have provided guidance for identifying patients and variants amenable to ASO intervention^6^, and patient-derived cellular models are accelerating empirical evaluation of personalized ASOs^7^. Yet identifying a pathogenic splicing defect does not specify where a therapeutic oligonucleotide should bind. For ultra-rare and patient-specific defects, this intervention-selection problem is especially consequential because only a small experimental panel and limited patient-derived material may be available.

Target-site selection therefore still often relies on oligonucleotide walks, in which overlapping ASOs are tiled across an exon and adjacent intronic sequence and promising windows are locally optimized by shifting the binding site or altering oligonucleotide length^8–10^. Although effective, this empirical search becomes costly when regulatory information is incomplete, relevant sites extend into introns, or candidate architectures become combinatorial. Dual-targeting ASOs, in which one oligonucleotide hybridizes to two separated pre-mRNA regions, have entered clinical testing in DMD^4,11,12^, but the resulting design space cannot be represented naturally by conventional scoring. The central computational problem is therefore not simply to predict splicing from sequence, but to rank where a therapeutic perturbation should be placed across heterogeneous sequence compartments and intervention architectures.

Recent approaches address complementary aspects of this problem. Machine-learning models can identify functional splice-switching oligonucleotide sites from splicing-regulatory features^13^, CRISPR– dCas13d screens can map proximal and distal regulatory elements at individual loci^14^, in silico deletion maps can nominate exonic regions for antisense modulation^15^, and KATMAP can infer cis-regulatory elements from splicing-factor perturbations^16^. Together, these studies show that diverse perturbations can reveal splicing regulation relevant to ASO design. However, the broader design problem remains unresolved: how to directly prioritize exon-skipping ASO target regions across exonic, intronic, exon–intron-spanning and split target architectures. We therefore formulated exon-skipping ASO design as region-level intervention ranking within a unified exon–intron sequence context.

Learning such a ranking is difficult because direct ASO activity data are sparse and heterogeneous. Single-nucleotide-variant (SNV)-based assays provide denser information about sequence positions at which local perturbation alters exon recognition. Although nucleotide substitution and ASO masking are mechanistically distinct, both identify splicing-sensitive sequence and may therefore provide complementary supervision for target-region prioritization.

Here we present eSkip2, an intervention-centric framework that jointly fine-tunes the genome-pretrained HyenaDNA model^17^ on experimentally measured ASO-induced exon skipping^9^ and SNV-derived exon-disruption profiles^18^.To support this joint learning, eSkip2 uses a common perturbation representation that encodes one or more ASO-binding regions or an SNV position within the sequence context. The same representation enables eSkip2 to rank exonic, intronic, exon–intron-spanning and split dual-targeting configurations within a common exon–intron context. eSkip2 also includes target-locus refinement without using experimental ASO outcomes from the locus being evaluated. Across held-out canonical exons and pseudoexons, eSkip2 prioritized active regions across multiple cell systems and chemistries. In a prospective DMD exon 46 study, the two most active new ASOs ranked within the top three and produced dose-dependent dystrophin restoration in patient-derived cells. Together, these results move genome sequence modeling from sequence-to-splicing prediction toward sequence-to-intervention ranking and illustrate a broader design principle for therapeutic sequence models under label scarcity: mechanistically related perturbation data can augment sparse direct intervention labels to prioritize actionable sites.

## Results

### eSkip2 prioritizes active regions across genes and contexts

eSkip2 receives an exon with 50-nt flanking intronic sequence on each side and assigns a score to one or more masked regions representing ASO binding (Fig. 1a). By modeling the perturbation within its local pre-mRNA context, the same representation accommodates exonic, intronic, exon–intron-spanning and split dual-targeting designs. We evaluated ranking with the area under the receiver operating characteristic curve (AUROC) because the intended use is to prioritize candidates rather than impose a universal activity threshold; threshold-dependent metrics are provided separately (Supplementary Table 1).

**Fig. 1.**
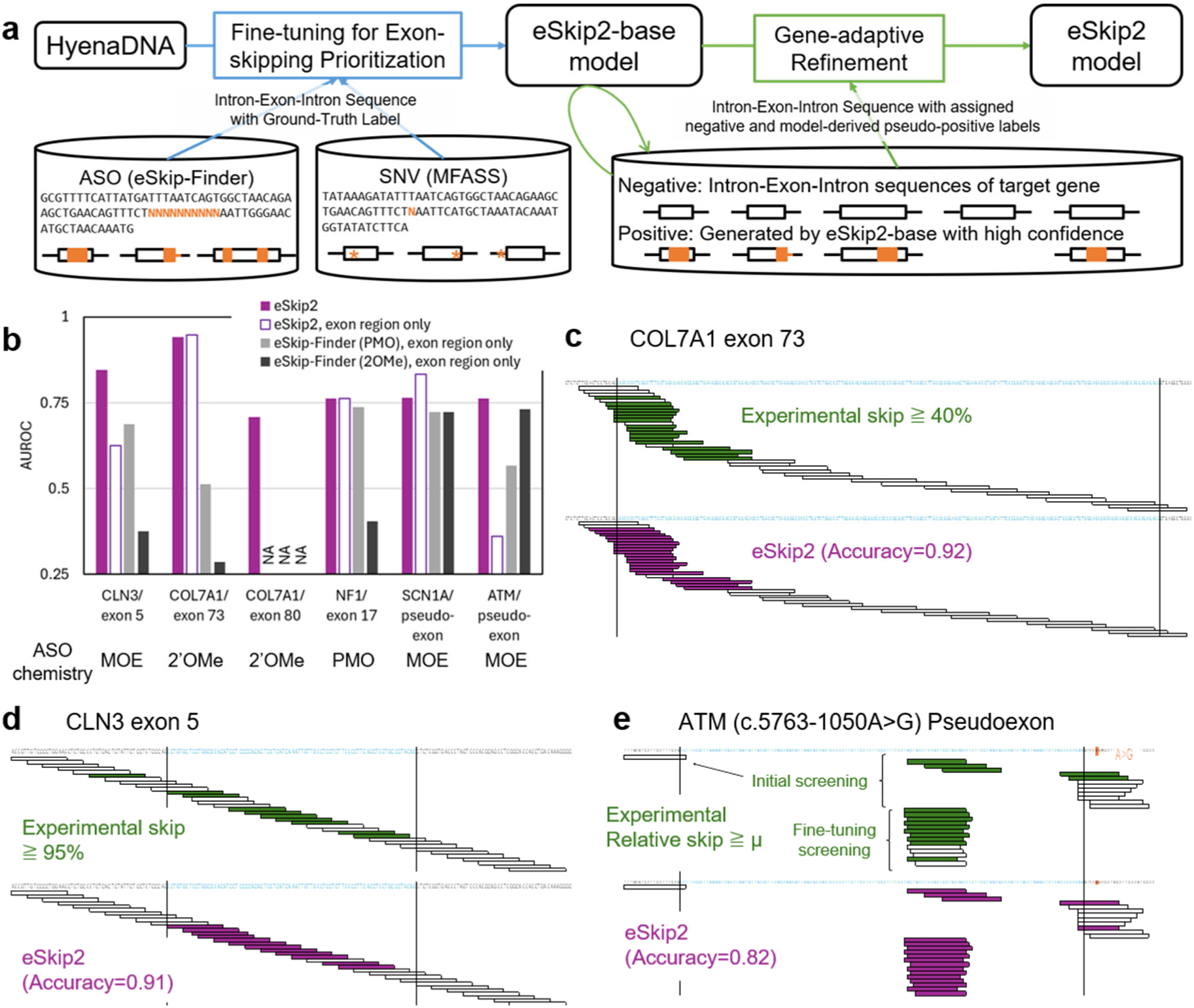
Cross-gene benchmarking of eSkip2 for ASO target-region prioritization. **a**, Overview of the eSkip2 framework. The pretrained HyenaDNA was fine-tuned for exon-skipping prioritization using labeled intron–exon–intron inputs derived from ASO-induced exon-skipping experiments and SNV-based exon-disruption data. Exons are shown as black rectangles and introns as lines; orange boxes and asterisks indicate ASO binding sites and SNVs, respectively. The cross-gene model (eSkip2-base) was then optionally refined for a target locus using a gene adaptation without target-specific experimental ASO activity labels. In this step, negatives were sampled from the target gene, and pseudo-positives were generated from the target gene sequence via model-guided masking; no experimental ASO activity labels from the evaluation target were used. Rounded rectangles indicate trained models, standard rectangles denote computational steps, and cylinders denote data sources. **b**, AUROC on benchmark test sets. AUROC was used as the primary benchmark summary because the task is target-region prioritization, and AUROC evaluates ranking without selecting a model-score cutoff once benchmark labels are fixed. Filled purple bars show eSkip2 performance across all available ASOs. For comparison with the previously reported eSkip-Finder models trained on PMO or 2′OMe ASO data, which are applicable only to exon-confined ASOs, eSkip2 AUROCs were recalculated on the corresponding exon-confined ASO subsets and are shown as purple-outlined open bars. The eSkip-Finder AUROCs calculated on these exon-confined subsets are shown in gray. For COL7A1 exon 80, the exon-confined subset contained only one ASO; therefore, exon-only AUROCs could not be calculated for either eSkip2 or eSkip-Finder and are shown as NA. Secondary descriptive metrics requiring a model-score cutoff are reported in Supplementary Table 1. **c**–**e**, Comparison of experimental exon-skipping outcomes (top; high-activity ASOs in green) and eSkip2 predictions (bottom; higher scores in purple) for representative benchmark targets. Each horizontal bar indicates one ASO binding site. Exonic and intronic sequences are shown in blue and black, respectively. Thresholded highlighting in these panels is shown for visualization only; benchmark comparisons rely on AUROC rather than on a single chosen model-score cutoff.

Across six benchmark targets held out from supervised ASO training and model-selection validation, eSkip2 prioritized active regions among all available exonic, intronic and exon–intron-spanning ASOs, with AUROCs of 0.71–0.94 (Fig. 1b and Supplementary Table 1). None of the five benchmark genes was included in the ASO-derived training set. Four were also absent from the SNV-derived MFASS data, and the one overlapping COL7A1 exon did not correspond to either evaluated COL7A1 exon (Supplementary Tables 2–4).

The benchmark panel mixed sparse literature-derived candidate sets with denser screens that more directly challenged region-level ranking; therefore, AUROC values should be interpreted in the context of dataset geometry (Supplementary Table 4). For CLN3, we evaluated both a stringent 95% skipping threshold and an alternative mean-based threshold to account for the unusually high activity of numerous exon-proximal ASOs^19^. eSkip2 achieved AUROCs of 0.85 and 0.77 under these definitions, respectively (Supplementary Table 1), arguing against dependence on a single favorable label definition.

The benchmark also tested transfer beyond model-development conditions. Evaluation datasets included MOE chemistry, non-RD cell systems including HeLa and HEK293 cells and patient-derived fibroblasts, and a wide concentration range (Fig. 1b and Supplementary Tables 4 and 5). Useful ranking across these conditions supports eSkip2 as a first-pass prioritization tool rather than a chemistry- or assay-specific predictor.

Region-level visualizations showed that eSkip2 recovered experimentally favored target zones across diverse splicing contexts (Fig. 1c–e). For COL7A1 exon 73, the model highlighted the 3′ splice site and adjacent exonic sequence while assigning lower scores to the exon center and 5′ end. For CLN3 exon 5, which has been extensively screened experimentally^19^, eSkip2 emphasized the central exonic region containing the strongest activity peak. In the ATM c.5763-1050A>G pseudoexon, a challenging ultra-rare context^6^, the model prioritized the 5′ splice site and exonic interior over the 3′ splice-site region, consistent with the experimentally productive design space. As expected from its emphasis on region prioritization, eSkip2 less effectively separated ASOs that differed by only one or two nucleotides within an already favorable region. This limitation was most visible in the ATM pseudoexon optimization set, where many closely related designs targeting the same active region received similarly high scores. We therefore interpret eSkip2 primarily as a search-space reduction tool that identifies promising regulatory regions, not as a final local optimizer for distinguishing near-identical oligonucleotides within an already favorable region.

To place the performance of eSkip2 in the context of previously reported prediction methods, we next compared it with eSkip-Finder, a machine-learning framework developed for exon-skipping ASO design. Because eSkip-Finder is applicable only to exon-confined ASOs, we separately recalculated eSkip2 AUROCs on the same exon-only candidate subsets (Fig. 1b). Across the common exon-confined subsets, eSkip2 had a median AUROC of 0.76, compared with 0.69 and 0.41 for the PMO- and 2′OMe-trained eSkip-Finder models, respectively, although performance varied among individual targets. Consistent with the local-resolution limitation described above, eSkip2 performed less well on the exon-confined ATM pseudoexon subset, which contained many closely overlapping ASOs targeting the same experimentally favored region (Fig. 1e). These results emphasize that the main advantage of eSkip2 is unified prioritization across exon– intron contexts rather than uniform superiority on every exon-only dataset.

The conventional eSkip-Finder (2′OMe) model^9^, which represents each ASO using manually defined sequence-derived and thermodynamic descriptors^20,21^, achieved an AUROC of 0.73 on the exon-confined ATM pseudoexon subset (Supplementary Figure 1). This result suggests that such predefined descriptors can still capture useful signals for local ASO prioritization^9,22^. This contrast reinforces the complementary roles of the two approaches: region-first prioritization by eSkip2, followed, where needed, by finer local optimization among closely related designs. GC count provides one example of a simple handcrafted feature that may be informative in specific contexts. Within the exon-confined ATM subset used for the eSkip-Finder comparison, GC count correlated with skipping activity (Supplementary Figure 2). However, when ASOs were tiled across BIM exon 3 and CLN3 exon 5 with minimal selection bias, GC count within the binding region showed no clear relationship with the experimental readout (Supplementary Figure 3). Thus, GC content alone cannot account for the activity patterns that eSkip2 seeks to capture, although its context-dependent association with activity suggests that explicit local sequence features may provide information complementary to the representations currently used by eSkip2.

### Prospective design identifies active dual-targeting ASOs

To test eSkip2 prospectively in a setting not addressed by existing tools, we applied it to dual-targeting ASOs for DMD exon 46, a locus not used for supervised model training or model-selection validation. Because general design rules for dual-targeting ASOs remain limited, we combined exhaustive eSkip2 scoring with a diversity-aware candidate-selection strategy. We enumerated all 24-mer candidates comprising two 12-nt complementary segments within the exon and its 50-nt flanks and scored them with the DMD-specific model (Fig. 2a,b). Because the highest-scoring coordinates formed a narrow cluster of similar designs, we combined model scores with local-maxima selection to preserve spatial and sequence diversity. Candidates whose two complementary segments merged into a continuous binding region were reclassified as single-targeting ASOs. Four additional single-targeting 24-mer ASOs were selected separately from the continuous single-targeting score profile (Fig. 2c). The final experimental panel therefore comprised 11 dual-targeting and 5 single-targeting 24-mer ASOs (Fig. 2d). Implementation details are provided in Methods.

**Fig. 2.**
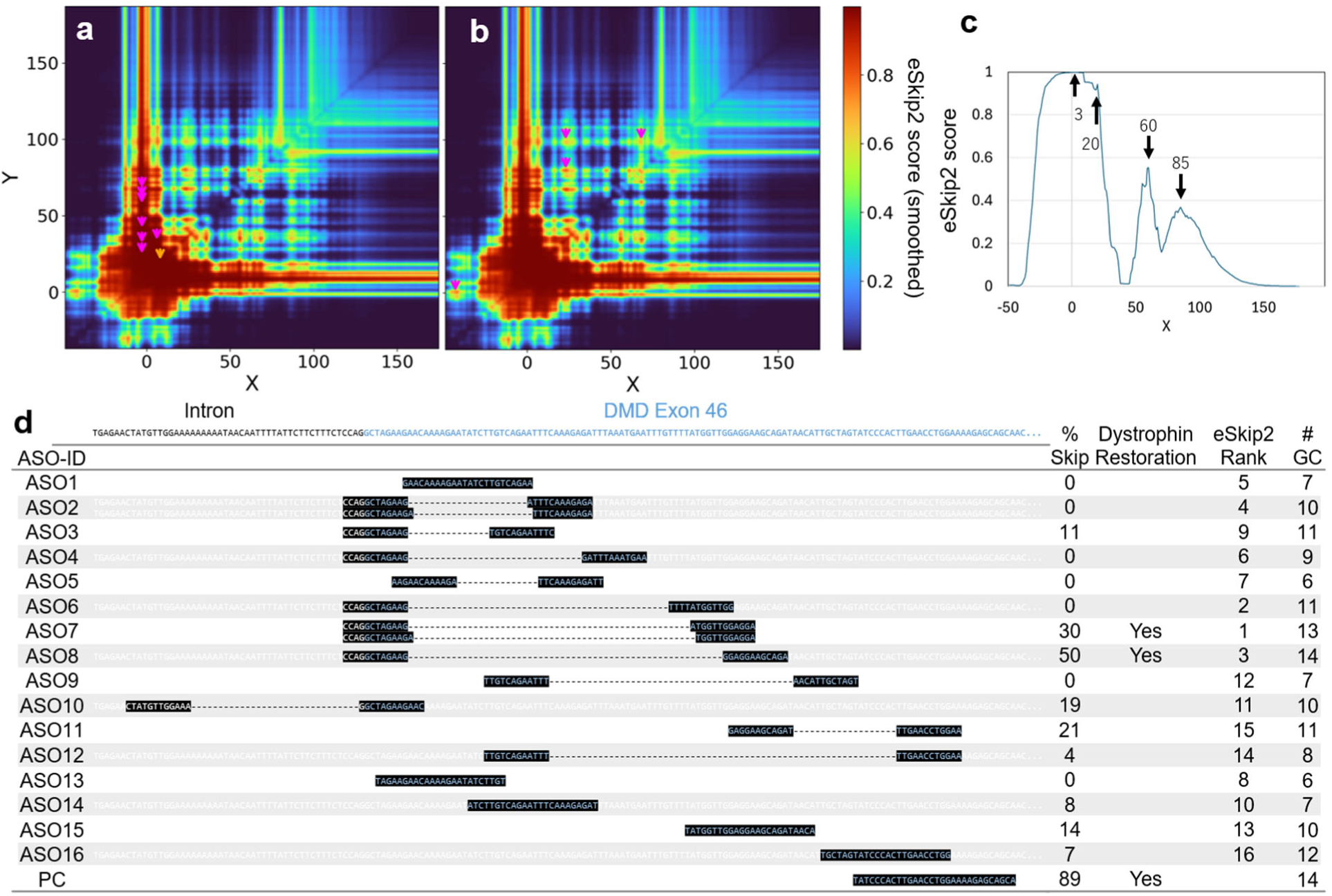
Design space and candidate selection for dual- and single-targeting ASOs at DMD exon 46. **a**, **b**, Two-dimensional heat maps of eSkip2 scores for dual-targeting ASOs enumerated across DMD exon 46 and its 50-nt flanking intronic sequence. Panels **a** and **b** correspond to local-maximum searches using 5 × 5 and 13 × 13 maximum-filter neighborhoods, respectively. The *x*- and *y*-axes denote the start positions of the first and second complementary segments, respectively, measured relative to the 3′ splice site. Magenta symbols indicate candidates selected as dual-targeting ASOs, while orange symbols indicate candidates initially identified in the dual-targeting search that were ultimately interpreted as continuous single-targeting designs because the two complementary segments merged into one binding region. **c**, eSkip2 scores for continuous single-targeting 24-mer ASOs tiled at a 1-nt resolution across the target region. The blue line shows the 10-nt moving average used to identify broad high-scoring regions for candidate selection. Arrows indicate the four single-targeting ASOs selected for experimental testing; the numbers denote the starting positions of the ASOs. **d**, Binding positions of the selected ASOs and their experimentally measured exon-skipping efficiencies following PMO treatment at 1 μM. Exonic sequence is shown in blue and intronic sequence in black. The positive control (PC) was the highest-activity exon 46-targeting PMO reported in the eSkip-Finder database and had a length of 30 nt. For ASO2 and ASO7, partial overlap between the two complementary segments allowed alternative interpretation as 13- and 11-nt segments; for downstream analyses, the higher of the corresponding eSkip2 scores was used, but this choice did not change the candidate rank order or the overall conclusions (Supplementary Table 6).

Separating model ranking from diversity-aware panel construction avoided an artificially favorable test consisting of near-duplicate ASOs. The prospective experiment therefore assessed whether eSkip2 could enrich active candidates while sampling distinct regions of the design landscape.

We tested the 16 candidates and one 30-mer positive control in MYOD1-converted urine-derived cells from three patients with DMD carrying an exon 45 deletion. At 1 μM, four dual-targeting ASOs and the positive control produced mean exon skipping of 19–50% (Figs. 2d and 3a; Supplementary Table 6). Sanger sequencing confirmed precise exon 46 skipping in treated cells, whereas untreated cells showed alternative splicing involving exon 44 (Fig. 3b,c).

**Fig. 3.**
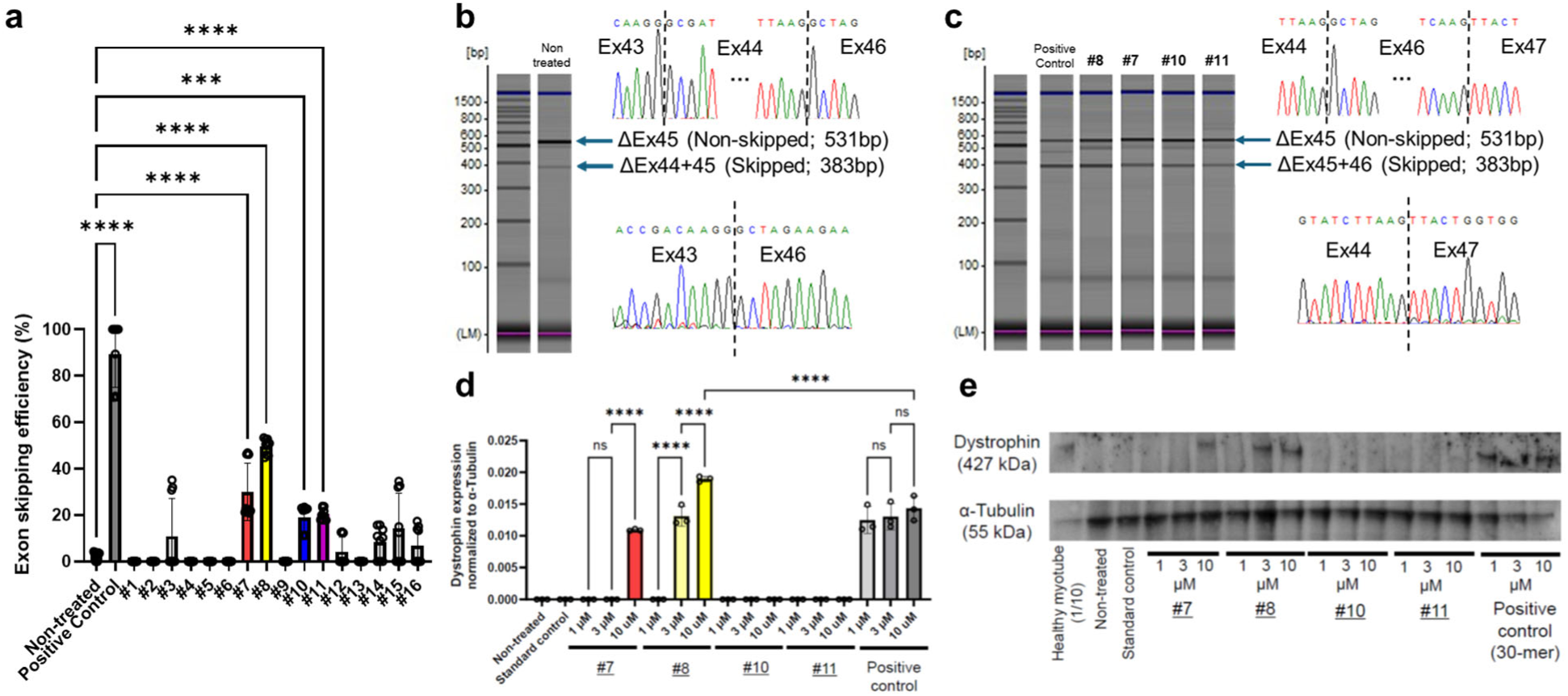
Exon 46 skipping and dystrophin restoration in *MYOD1*-converted urine-derived cells from patients with DMD carrying an exon 45 deletion. **a**, Exon 46 skipping efficiencies in *MYOD1*-converted urine-derived cells from three patients with DMD carrying an exon 45 deletion after treatment with 11 dual-targeting 24-mer ASOs, 5 single-targeting 24-mer ASOs, and 1 single-targeting positive-control 30-mer at 1 μM during myogenic differentiation. Data are mean ± s.d.; \*\*\**P* < 0.001, \*\*\*\**P* < 0.0001. One-way ANOVA with Sidak’s multiple-comparisons test was used. **b**, RT-PCR and sequencing showing alternative splicing involving exon 44 in untreated cells. **c**, Representative RT-PCR products and Sanger sequencing chromatograms confirming precise exon 46 skipping (ΔEx45–46; 383 bp) relative to the non-skipped transcript (531 bp). **d,e**, Immunoblot analysis of dystrophin (427 kDa) and α-tubulin (55 kDa) in *MYOD1*-converted urine-derived cells from three patients with DMD carrying an exon 45 deletion after treatment at 1, 3, and 10 μM with ASOs #7, #8, #10, and #11 and the positive control. Fifteen micrograms of total protein were loaded per lane (1.5 μg for healthy myotubes). Quantification was normalized to α-tubulin and referenced to healthy myotubes. Data are mean ± s.d.; \*\*\*\**P* < 0.0001. Two-way ANOVA with Sidak’s multiple-comparisons test was used.

ASO8 and ASO7 were the two most active new candidates, producing mean exon-skipping efficiencies of 50% and 30%, respectively. eSkip2 ranked ASO7 first and ASO8 third. Under random ordering of the 16 prospectively tested candidates, the probability that the two most active ASOs would both appear in the top three is 0.025. Thus, within the 16-candidate prospectively tested panel sampled from the combinatorial design space, the strongest activities were concentrated near the top of the eSkip2 ranking.

Protein-level analysis supported the functional relevance of this enrichment. ASO7 and ASO8 produced dose-dependent dystrophin restoration at 1, 3, and 10 μM (Fig. 3d,e). Although the positive-control ASO produced detectable dystrophin at lower concentrations, ASO8 showed the strongest restoration among the newly designed candidates at 10 μM. By contrast, separate analysis of the remaining 12 ASOs not included in Fig. 3d,e—all of which produced <20% exon skipping at 1 μM—showed no detectable dystrophin restoration at 10 μM (Supplementary Figure 4).

The two most active ASOs also had high GC counts (Fig. 2d), but GC content alone did not reproduce activity patterns in tiled BIM and CLN3 screens (Supplementary Figure 3) and was not used as a simple selection filter. The prospective DMD experiment therefore supports the practical use of joint exon–intron ranking to identify functional candidates in a dual-targeting design space.

### Joint ASO and SNV training supports gene adaptation

We next assessed the contributions of joint ASO–SNV training and gene-adaptive refinement using ablated models that lacked gene adaptation, ASO data or SNV data (Fig. 4a and Supplementary Figures 5–8). These analyses were performed after the fixed benchmark and prospective experiment and did not affect the model or candidates reported in Figs. 1–3.

**Fig. 4.**
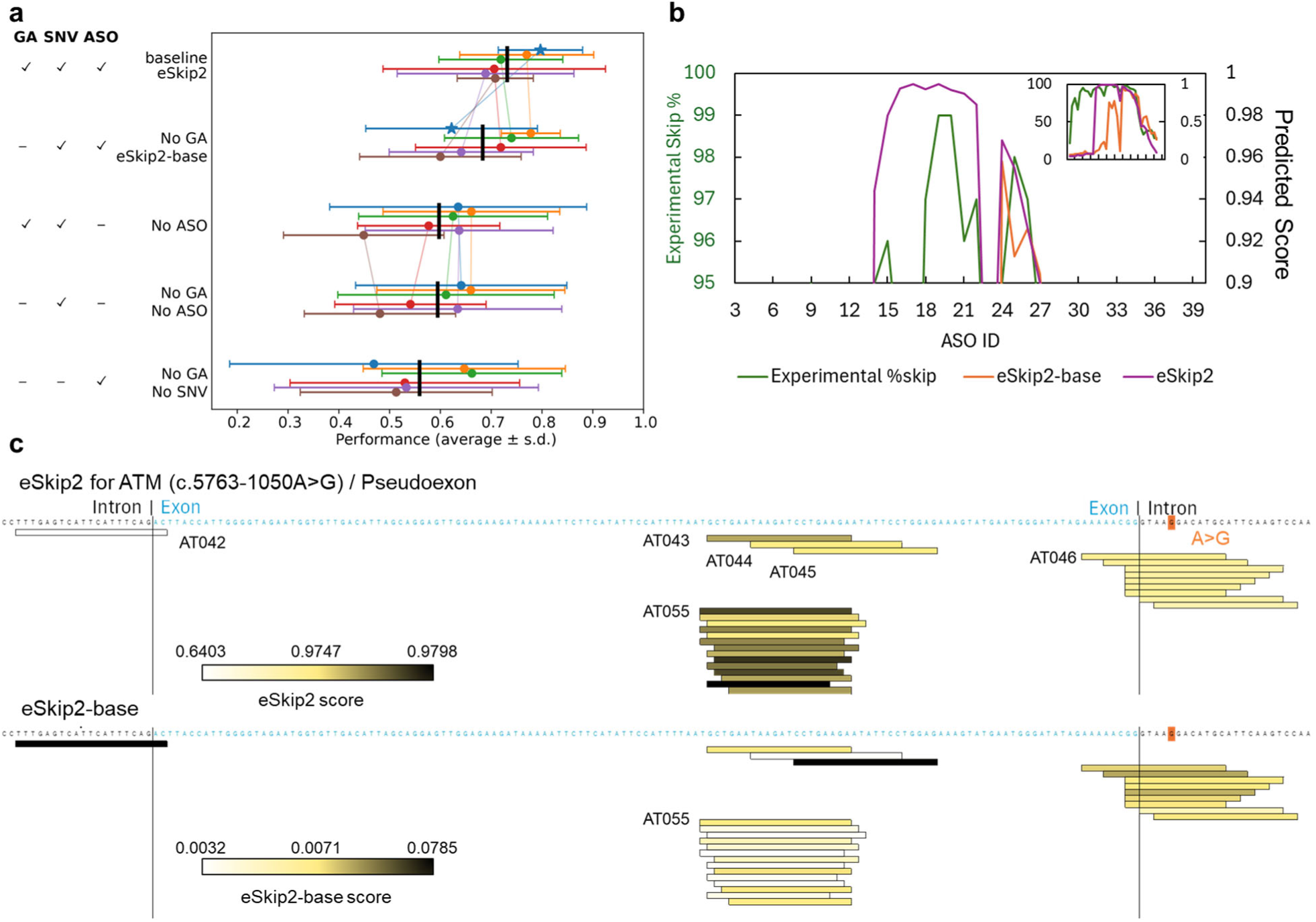
Effects of joint ASO–SNV training and ASO-label-free gene-adaptive refinement. a, Ablation analysis across model configurations differing in the availability of gene-adaptive refinement (GA), SNV-derived data and ASO-derived data. Each colored point represents one of the top five validation-ranked random-seed runs within the corresponding model class. Each point shows the mean AUROC across evaluation targets, and the horizontal bar indicates the corresponding standard deviation across targets; black bars indicate the mean across the five displayed seeds. Because each gene-adapted model was initialized from the corresponding base model trained with the same seed, thin lines connect paired runs. Stars mark the pre-specified validation-selected runs from the original hyperparameter searches. The 50-seed sweep was performed after the fixed benchmark and prospective analyses and was used only for post hoc robustness analysis. **b**, Experimental exon-skipping efficiencies (left axis, green) and model-predicted scores (right axis) for CLN3 exon 5 across individual ASOs. Predictions from eSkip2 are shown in purple and those from eSkip2-base in orange. The inset summarizes the distribution of predicted scores across all evaluated ASOs. Curves correspond to the pre-specified validation-selected run indicated by the star in **a**. **c**, Predicted scores for ASOs targeting the ATM pseudoexon (c.5763-1050A>G) generated by eSkip2 (top) and eSkip2-base (bottom). Each horizontal bar represents one ASO binding site and is colored according to the relative predicted score. The orange marker indicates the pathogenic A>G variant responsible for pseudoexon activation. Exonic sequence is shown in blue and intronic sequence in black. Predictions correspond to the pre-specified validation-selected run indicated by the star in **a**.

We first selected the hyperparameter setting for each pre-adaptation training scheme through the analyses shown in Supplementary Figures 5–7. After fixing these settings, we independently trained 50 models with different random seeds for each scheme to assess seed-to-seed variability (Supplementary Figure 8).

Joint training yielded more models with simultaneously high AUROC on BIM exon 3 and DMD exon 44 than either single-source training scheme (Supplementary Figure 8). This pattern suggests that the main advantage of joint training was a greater likelihood of obtaining models that performed well across heterogeneous validation targets, rather than a gain specific to either target.

For each scheme, the five highest-ranked seeds were selected, highlighted in Supplementary Figure 8, and used as starting models for the gene-adaptive refinement evaluated in Fig. 4a. Comparisons of each selected seed before and after gene adaptation showed larger gains when the pre-adaptation model assigned low scores to experimentally supported target regions. In contrast, gene adaptation had little effect when these regions were already highly ranked. This behavior is consistent with target-specific correction rather than uniform improvement across targets.

At CLN3 exon 5, the gene-adaptive refinement increased AUROC from 0.71 to 0.85 and recovered all three experimentally supported high-activity regions (Fig. 4b). At the ATM pseudoexon, it reduced emphasis on the weakly active 3′ splice-site region and shifted scores toward the causative-variant and central pseudoexonic regions, including the region targeted by the most active experimentally identified ASO, increasing AUROC from 0.36 to 0.76 (Fig. 4c).

These results support complementary roles for the two stages: ASO and SNV perturbations improve the robustness of the cross-gene model, whereas ASO-label-free gene adaptation can rescue locus-specific failures without using experimental ASO labels from the target.

## Discussion

eSkip2 addresses a practical bottleneck in exon-skipping therapy development: deciding which pre-mRNA regions warrant experimental testing. Its unified exon–intron representation accommodates exonic, intronic, exon–intron-spanning and split dual-targeting designs, while the masked-region formulation focuses the model on the local perturbation produced by ASO binding. Accordingly, eSkip2 serves as an upstream region-prioritization tool that systematically narrows the candidate space before ASO synthesis and experimental testing.

Three findings support this role. First, eSkip2 generalized to held-out canonical exons and pseudoexons across multiple chemistries and cellular systems. Second, joint learning from ASO and SNV perturbations improved robustness under limited ASO supervision. Third, prospective application to DMD exon 46 placed the two most active new ASOs within the top three ranks and linked model-guided prioritization to dose-dependent dystrophin restoration. The prospective result is particularly important because dual-targeting designs expand combinatorially and only a small fraction can be tested.

Benchmark performance should therefore be interpreted as evidence that eSkip2 can prioritize promising target regions, rather than as evidence of universally calibrated activity prediction. The evaluation datasets differ in ASO chemistry, concentration, cell type, assay readout, and label definition, and several consist of literature-derived candidate sets rather than unbiased saturation screens. Consequently, a single absolute activity threshold cannot be meaningfully applied across all benchmarks. At the same time, this heterogeneity provides a realistic test of whether eSkip2 can consistently rank more active ASOs above less active ones across diverse experimental settings. The enrichment of the two most active ASOs among the top three ranked candidates in the prospective DMD exon 46 experiment provides independent support for this intended use.

eSkip2 operates at two levels. eSkip2-base learns a cross-gene prior over splicing-sensitive regions from ASO masking and SNV perturbations. Gene-adaptive refinement then tailors this prior using only target-locus sequence and model-derived pseudo-labels. This separation is valuable for rare and individualized targets, where the need for computational guidance is greatest precisely because labeled ASO data do not yet exist.

The complementary contribution of ASO and SNV data is biologically plausible. SNV assays densely sample sequence positions that influence exon recognition, whereas ASO experiments connect these regulatory signals to the intervention of interest—masking a region to induce skipping. The ablation results indicate that neither source is fully interchangeable with the other. Gene adaptation, in contrast, is best viewed as a conditional rescue mechanism for atypical loci such as pseudoexons, not as a uniformly beneficial additional training stage.

Several limitations define the current scope. eSkip2 ranks regions but does not reliably distinguish one-nucleotide shifts or small length changes within the same active window. It does not explicitly encode ASO chemistry, duplex thermodynamics, delivery, off-target binding or downstream safety, and it was trained with binary rather than continuous activity labels. The prospective study tested one DMD exon in patient-derived cells, so additional prospective studies across genes, chemistries and delivery systems will be needed to establish the breadth of generalization.

These limitations suggest a modular development path in which eSkip2 is used to nominate regions, followed by chemistry-aware local optimization and experimental triage. Continuous-response learning, explicit oligonucleotide features and broader prospective datasets should improve resolution among closely related candidates. The benchmark-ready multi-gene resource and public implementation provided here enable such comparisons. More broadly, the study shows how genome-pretrained models and related perturbation data can be combined to support therapeutic design when direct labels are scarce.

## Methods

### Training, validation and benchmark datasets

To develop a model for ASO target-region prioritization, we assembled training data from two experimentally grounded sources of splicing perturbation: (i) ASO-induced exon-skipping experiments, where *in vitro* assays measured the presence or absence of skipping after oligonucleotide treatment^9^, and (ii) SNV-induced exon-disruption data from the MFASS dataset, a massively parallel assay that quantifies the effect of SNVs on exon recognition^23^. The rationale for combining these sources is that both perturbations identify regulatory regions whose local alteration can change exon inclusion, even though the molecular mechanisms differ.

For ASO training data, we extracted experimentally curated records from the eSkip-Finder database. To reduce technical heterogeneity during model development, we restricted the training set to experiments satisfying the following criteria:

1. Only experiments using human RD cells were included.
2. Consequently, ASO delivery methods were restricted to electroporation or transfection.
3. ASO chemistries were limited to PMO and 2′OMe. These chemistries are the most extensively represented among experimentally curated exon-skipping datasets and are also used clinically. Although eSkip-Finder contains additional chemistries such as MOE and locked nucleic acids, direct pooling of these data would have introduced larger technical heterogeneity.
4. ASO concentrations were restricted to 10–12.5 μM for PMO and 0.1 μM for 2′OMe. In our source data, the exon-skipping effects of PMO at ∼10 μM were broadly comparable to those of 2′OMe at 0.03–0.1 μM.

Exon-skipping efficiency was defined as skipped mRNA/(skipped mRNA + non-skipped mRNA). Samples with ≥40% skipping were labeled positive and those with <40% skipping were labeled negative. Samples with contradictory or duplicate labels under this definition were excluded. The final ASO training set comprised six DMD exons and one MSTN exon (Supplementary Table 2). Sequences were obtained from NCBI Reference Sequences NG_012232 (DMD) and NG_009800 (MSTN).

For ASO data, each model input consisted of the target exon plus 50 nt of flanking intronic sequence on each side. To represent ASO binding, the complementary RNA region was masked with the N token, which HyenaDNA interprets as an unknown nucleotide. We adopted this representation to approximate the local occlusion created by ASO hybridization while preserving sequence length and positional continuity, thereby enabling a unified encoding of single- and dual-targeting designs.

SNV inputs were derived from MFASS sequences, which contain ∼40 nt of upstream and ∼30 nt of downstream intronic context. The SNV position was replaced with the N token. When different substitutions at the same site produced conflicting outcomes, the site was labeled positive if any substitution induced exon deletion.

We used two independent validation datasets, DMD exon 44 and BIM exon 3, to select models with good cross-context performance (Supplementary Table 3). DMD exon 44 included ASOs targeting the 3′ splice site and dual-targeting designs. BIM exon 3 used fixed-length 18-nt ASOs and broad tiling across exon-proximal sequences (Supplementary Figure 9)^24^. Because the BIM assay quantified the abundance of the retained exon rather than a direct skipping percentage, samples were labeled positive when the E3 ratio fell below the dataset mean minus half the standard deviation; ambiguous samples near the threshold were excluded.

For final benchmarking, we used 6 evaluation targets, each with at least 10 tested ASOs (Supplementary Table 4): 2 COL7A1 exons, CLN3 exon 5 under 2 operational label definitions, the SCN1A pseudoexon, the ATM c.5763-1050A>G pseudoexon, and NF1 exon 17^6,19,25–28^. These loci were absent from the ASO-derived training set. To assess potential overlap with MFASS, we intersected benchmark coordinates with GENCODE v27 annotations. Four of the five benchmark genes were absent from MFASS training data, and the single overlapping COL7A1 exon in MFASS did not correspond to the benchmark exons analyzed here.

Operational positive/negative thresholds were defined separately for each benchmark to derive binary labels from heterogeneous experimental readouts (Supplementary Table 4). In most datasets, ASOs were labeled positive when their exon-skipping efficiency was equal to or greater than the mean activity within that dataset. Exceptions were made for datasets reported only as categorical activity ranges, for which an approximate threshold was derived from the reported categories, and for CLN3 exon 5, for which a stringent 95% skipping threshold was used for the primary benchmark to avoid labeling nearly all exon-proximal ASOs as positive. A mean-based CLN3 threshold was also evaluated as a sensitivity analysis. These thresholds were used only to derive binary benchmark labels. AUROC was then calculated without choosing a model-score cutoff, whereas secondary descriptive metrics such as Matthews correlation coefficient (MCC), accuracy, and precision additionally required a dataset-specific score cutoff. For these descriptive summaries, the score cutoff was selected within each dataset to maximize MCC and was not used for model selection or for the primary benchmark comparison.

All experiments except those involving SCN1A used electroporation or transfection for cellular ASO delivery, which reduces the likelihood of sequence-specific uptake dominating the benchmark results. We therefore did not explicitly model cellular entry. The experimental conditions underlying each dataset are summarized in Supplementary Table 5.

### Construction of the eSkip2-base model

Publicly available, well-curated ASO skipping datasets remain limited in scale, making *de novo* training of a high-capacity model difficult. We therefore used transfer learning from a genome foundation model. Among recent sequence models such as DNABERT^29^ and HyenaDNA^17^, HyenaDNA offers a favorable balance between performance and computational cost on genomic tasks^30^. We selected the pretrained eight-layer 160k-context variant (hyenadna-medium-160k-seqlen) for downstream fine-tuning.

We formulated exon-skipping prioritization as a binary classification task where the model predicts whether a masked sequence corresponds to a perturbation that induces exon skipping. Training used 568 ASO examples curated from eSkip-Finder and 30,015 SNV examples from MFASS. Because the combined training set was highly imbalanced (1,295 positives and 29,288 negatives), we used a class-weighted loss with a 23-fold weight on positive samples.

We optimized learning rate, dropout, number of layers retained from the pretrained model, batch size, and number of epochs over the ranges listed in Supplementary Table 7. Because the transferability of pretrained representations varies across network depth and target tasks^31^, we included the number of retained pretrained layers as a search dimension. Candidate models were evaluated on the DMD exon 44 and BIM exon 3 validation sets, and hyperparameter settings were ranked by the sum of Z-score-normalized AUROCs across the two validation tasks (Supplementary Figure 5). GS_231, the single run selected using this validation-only criterion, was fixed before prospective candidate selection for DMD exon 46. The selected configuration is listed in Supplementary Table 8. Training with this setting required approximately 30 min on an NVIDIA A100 40 GB GPU (NVIDIA, Santa Clara, CA, USA).

Within the selected hyperparameter configuration, predictions were stabilized by averaging model weights over the final 10% of epochs. The resulting cross-gene model was designated eSkip2-base. The fixed GS_231 eSkip2-base model was used to initialize each target-specific gene-adapted model. Unless otherwise noted, Fig. 1 and Supplementary Table 1 report predictions from the corresponding gene-adapted eSkip2 model. No experimental ASO activity labels from the evaluation target were used during the gene-adaptive refinement. The later 50-seed analyses in Fig. 4a and Supplementary Fig. 8 were performed separately and did not influence the fixed benchmark results or prospective candidate selection.

### Gene-adaptive refinement

Gene adaptation was designed to reflect practical deployment: the genomic sequence of the target locus was known, but experimentally measured ASO outcomes for that target were unavailable. No experimental ASO activity labels from the evaluation target were used during this refinement step.

For a given target gene, we collected constitutive exons and their flanking intronic sequence and labeled them as negatives. The exon sets and RefSeq accessions used for gene-adaptive refinement are listed in Supplementary Table 9. Pseudo-positives were then generated using model-guided *in silico* masking. We first excluded negative sequences that the current model already scored >0.5, because these were likely difficult or ambiguous negatives. For each remaining sequence, all possible contiguous masked segments of 20 nt were evaluated, or 18 nt for targets whose experimental screens used fixed 18-mers (CLN3, ATM, and SCN1A). The highest-scoring masked sequence for a given exon was retained as a pseudo-positive if its score exceeded 0.5.

The intended target exon or pseudoexon was included in the refinement set only through this sequence-only pseudo-labeling procedure. No experimentally observed skipping outcomes from the evaluation target were introduced. Pseudo-positives were split approximately evenly between refinement-training and refinement-validation sets, and the corresponding negative sequences were partitioned in parallel.

We illustrate the procedure using COL7A1 exon 73. The COL7A1 genomic reference (NG_007065.1) contains 118 exons; exons 1 and 118 were excluded because one flanking intron is absent. The remaining 116 exons were represented with 50-nt flanks and used as negatives. Eleven sequences with base-model scores >0.5 were removed from pseudo-label generation, leaving 105 candidates. Exhaustive masking of these sequences identified 97 pseudo-positives with scores >0.5, including exon 73. The final refinement dataset contained 49 pseudo-positives and 68 negatives for training, and 48 pseudo-positives and 48 negatives for validation.

For special cases such as the ATM c.5763-1050A>G pseudoexon, the mutant sequence containing the pseudoexon plus 50-nt flanks was treated as a candidate for pseudo-labeling and also as a negative example, because pseudoexon inclusion represents the opposite of the desired skipping outcome. The corresponding wild-type sequence, which lacks pseudoexon inclusion, was treated as a true positive.

Fine-tuning started from eSkip2-base using a small learning rate (Supplementary Table 10). The model with the lowest validation loss was retained as the gene-adapted model, referred to as eSkip2. Even for large genes, computation remained modest: for COL7A1, pseudo-label generation for 18,129 masked sequences required approximately 12 min, and subsequent fine-tuning on 213 total sequences required approximately 3 min on an NVIDIA A100 40 GB GPU. Model inference required well below 1 s per sequence, facilitating scalable *in silico* screening.

### Design of dual- and single-targeting ASOs using eSkip2

Dual-targeting ASOs were modeled as two complementary 12-nt binding segments within the DMD exon 46 target sequence and its 50-nt flanking intronic sequence. Coordinates were defined relative to the 3′ splice site, which was assigned position 0. For every possible pair of start positions, we generated the corresponding masked sequence representation and scored it with the DMD-specific eSkip2 model. This exhaustive enumeration produced a two-dimensional prediction table that was visualized as a heat map (Fig. 2a, b).

To facilitate candidate selection, the score matrix was smoothed with a Gaussian filter implemented in SciPy^32^. Local maxima were then identified with a square maximum filter using either a 5 × 5 or a 13 × 13 neighborhood. We additionally required candidate peaks to show local plateau-like behavior, defined as similar values across the eight neighboring cells within a 20% relative tolerance, to suppress isolated noisy maxima. Designs in which the two segments were separated by only 1–9 nt were not carried forward in the final candidate panel. Application of the 5 × 5 neighborhood yielded eight candidates (ASO1–ASO8). The 13 × 13 neighborhood was subsequently used to identify four additional candidates after duplicate and near-duplicate designs had been removed (ASO9–ASO12). Candidates in which the two complementary segments were directly contiguous were reclassified as continuous single-targeting 24-mer ASOs.

In parallel, continuous single-targeting 24-mer ASOs were scored at 1-nt resolution across the same target region. A 10-nt moving average of these scores was used to identify broad high-scoring regions and guided the selection of four additional single-targeting ASOs (ASO13– ASO16). Finally, the eSkip2-designed candidate panel comprised 11 dual-targeting and 5 single-targeting 24-mer ASOs. A previously reported single-targeting 30-mer ASO was included separately as a positive control.

### Ablation analysis and seed selection

To dissect the roles of SNV-derived data, ASO-derived data, and gene adaptation, we trained ablated variants using only SNV data (No ASO), only ASO data (No SNV), or the full training scheme without gene adaptation (eSkip2-base). Hyperparameters for the SNV-only and ASO-only models were optimized using the same validation procedure used for the full model (Supplementary Figures 6 and 7; Supplementary Tables 11 and 12).

After selecting one hyperparameter configuration for each model class, we trained 50 independent random-seed replicates to assess stochastic variability. For each replicate, we computed an unweighted average of the model parameters across checkpoints from the final 10% of training epochs, using the same checkpoint-averaging procedure applied to the fixed eSkip2-base model. These seed sweeps were performed after the fixed benchmark and prospective analyses and were used only to characterize robustness; they did not redefine the models reported in Fig. 1 or the candidate ranking used in Figs. 2 and 3. For compact visualization in Fig. 4a, we highlighted the top five runs within each model class according to the validation ranking, whereas the full distributions across all 50 seeds are shown in Supplementary Fig. 8. Gene-adaptive refinement was also attempted for the ASO-only configuration, but pseudo-label generation yielded too few positive examples for stable refinement; therefore, this configuration was not pursued further.

## Experimental validation of eSkip2-designed ASOs

### Cell culture

UDCs were isolated from the urine of each patient as described previously^33,34^. Briefly, urine samples were centrifuged at 400 × *g* for 10 min at 15–25 °C. The cell pellet was resuspended in growth medium comprising a 1:1 mixture of high-glucose Dulbecco’s modified Eagle medium and REGM Bullet Kit (Lonza, Basel, Switzerland), supplemented with 15% tetracycline-free fetal bovine serum, 0.5% GlutaMAX-I (Gibco; Thermo Fisher Scientific, Waltham, MA, USA), 0.5% non-essential amino acids (Thermo Fisher Scientific), basic fibroblast growth factor (Sigma-Aldrich, St. Louis, MO, USA), platelet-derived growth factor-AB (Peprotech, Cranbury, NJ, USA), epidermal growth factor (Peprotech), amphotericin B, and penicillin/streptomycin. Thereafter, cells were seeded in gelatin-coated plates and cultured at 37 °C under 5% CO_2_ for 3 days. Medium was replaced on day 4 and every other day thereafter. Mycoplasma contamination was monitored routinely with the TaKaRa PCR Mycoplasma Detection Set (Takara Bio, Shiga, Japan). CD90-positive UDCs were isolated using fluorescein isothiocyanate-conjugated anti-human CD90 antibody (#328107; BioLegend, San Diego, CA, USA) and sorted on a SONY FACS SH800S instrument (Sony Biotechnology, San Jose, CA, USA). For direct reprogramming into myotubes, UDCs were transduced with a doxycycline-inducible *MYOD1* retroviral vector at a multiplicity of infection of 200 in 60-mm gelatin-coated plates^33,34^. MYOD1-UDCs were differentiated into myotubes by switching to differentiation medium containing doxycycline. Medium was refreshed every 3 days.

### ASO synthesis and transfection

Newly designed ASOs targeting DMD exon 46 were synthesized by Gene Tools (Philomath, OR, USA). ASOs were introduced into MYOD1-UDCs from patients with DMD on day 7 of differentiation using Endo-Porter (Gene Tools) at final concentrations of 1, 3, or 10 μM. After 72 h of incubation, cells were transferred to fresh ASO-free differentiation medium and analyzed on day 14.

### RNA analysis of exon skipping efficiency

Total RNA was extracted from UDCs and MYOD1-UDCs using an RNeasy kit (Qiagen, Hilden, Germany). Exon 46 skipping was quantified using a one-step RT-PCR reaction (QIAGEN) under the following cycling conditions: 50 °C for 30 min; 95 °C for 15 min; 35 cycles of 94 °C for 1 min, 60 °C for 1 min, and 72 °C for 1 min; and 72 °C for 7 min. Products were analyzed on a MultiNA microchip electrophoresis system (Shimadzu, Kyoto, Japan). Exon-skipping efficiency (%) was calculated as exon 45–46-deleted PCR fragment / (exon 45–46-deleted PCR fragment + exon 45-deleted PCR fragment) × 100. Primer sequences were as follows: forward (5′→3′), ACAACAAAGCTCAGGTCGGA; and reverse (5′→3′), CACTTACAAGCACGGGTCCT.

### Sanger sequencing

RT-PCR products from untreated and ASO-treated MYOD1-UDCs were purified using a Wizard SV Gel and PCR Clean-Up System (Promega, Madison, WI, USA) and subjected to Sanger sequencing by Eurofins Japan (Tokyo, Japan).

### Dystrophin immunoblotting

Total protein was extracted from cultured cells in RIPA buffer supplemented with cOmplete Mini Protease Inhibitor Cocktail (Roche, Basel, Switzerland). Afterward, lysates were sonicated using 10 pulses of 10 s in cold water with a SONIFIER 250D (Branson Ultrasonics, Brookfield, CT, USA) and centrifuged at 15,000 × *g* for 15 min at 4 °C. Protein concentration was measured using a BCA assay (Thermo Fisher Scientific). Samples were mixed with NuPAGE LDS Sample Buffer (Thermo Fisher Scientific), denatured at 70 °C for 10 min, and separated on NuPAGE Novex Tris–Acetate 3–8% gels (Invitrogen; Thermo Fisher Scientific) at 150 V for 70 min before transfer to PVDF membranes. Next, membranes were blocked with 5% ECL Prime blocking agent (Cytiva, Marlborough, MA, USA) in PBS containing 0.1% Tween 20 for 1 h at 15–25 °C and incubated overnight at 4 °C with anti-dystrophin (1:20; NCL-DYS1; Leica Biosystems, Wetzlar, Germany) and anti-α-tubulin (1:1,200; T5168; Sigma-Aldrich). After washing, membranes were incubated with anti-mouse horseradish peroxidase-conjugated secondary antibodies (1:10,000; NA9310; Cytiva) for 1 h at 15–25 °C. Subsequently, signals were detected using ECL Prime Western Blotting Detection Reagent (GE Healthcare, Chicago, IL, USA) on a ChemiDoc MP imaging system (Bio-Rad Laboratories, Hercules, CA, USA) and quantified using Image Lab software (Bio-Rad Laboratories).

### Statistical analysis

Statistical analyses were performed in GraphPad Prism v9.2.0 (GraphPad Software, La Jolla, CA, USA). Quantitative data in Fig. 3 are presented as mean ± s.d. Comparisons in Fig. 3a and Fig. 3d were performed using one-way and two-way ANOVA, respectively, followed by Sidak’s multiple-comparisons test. *P* < 0.05 was considered statistically significant.

## Use of generative AI tools

A large language model was used to assist manuscript revision and language editing. It was not used to generate data, conduct analyses or determine scientific conclusions. All suggested text was reviewed and approved by the authors.

## Data availability

The curated ASO activity data underlying the training, validation and benchmark sets, including all benchmark splits, will be released through the eSkip-data repository (https://github.com/riken-yokohama-AI-drug/eskip-data) upon acceptance or upon publication. The complete data are available to editors and reviewers during peer review.

## Code availability

Code for reproducing the benchmark analyses and gene-adaptive refinement for DMD exon 46, together with trained model weights, configuration files and environment specifications, is publicly available at https://github.com/riken-yokohama-AI-drug/eskip2.

## Ethics approval and informed consent

This study was approved by the Ethics Committee of the National Center of Neurology and Psychiatry (approval ID: A2018-029), and written informed consent was obtained from all patients with DMD carrying an exon 45 deletion before urine collection. All experiments were conducted in accordance with the relevant guidelines and regulations.

## Acknowledgements

This research was supported by Grants-in-Aid for Research on Nervous and Mental Disorders [grant number 5-7] and by the Japan Agency for Medical Research and Development (AMED) under grant number 25nk0101112s0501. Computational resources were provided by the Hokusai Big Waterfall2 and RAIDEN supercomputers. We thank Dr Tomoharu Isobe for assistance with organizing the GitHub repository.

## Author contributions

S.C.: Conceptualization, Methodology, Software, Formal analysis, Investigation, Data curation, Writing – original draft, Visualization, Project administration, Funding acquisition.

K.K.: Formal analysis, Investigation, Resources, Writing – original draft, Writing – review & editing, Visualization.

S.S.: Investigation, Resources, Writing – review & editing.

U.S.H.: Data curation, Writing – review & editing.

H.W.-C.: Data curation, Writing – review & editing.

M.N.A.S.: Data curation, Writing – review & editing.

J.L.: Data curation, Writing – review & editing.

K.M.: Data curation, Writing – review & editing.

F.U.-O.: Methodology, Writing – review & editing.

T.Y.: Validation, Resources, Writing – review & editing, Supervision, Funding acquisition.

Y.A.: Resources, Writing – review & editing, Supervision, Funding acquisition.

Y.O.: Conceptualization, Methodology, Resources, Writing – review & editing, Supervision, Project administration, Funding acquisition.

