## Supplemental file for "eSkip2 prioritizes exon-skipping antisense oligonucleotide target regions across exon–intron contexts"

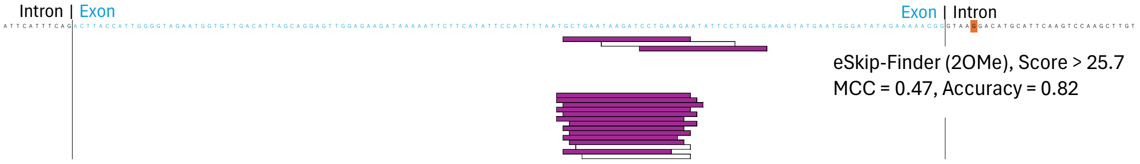


**Supplementary Figure 1 | eSkip-Finder predictions for exon-confined ASOs in the ATM pseudoexon.**

ASOs targeting the ATM c.5763–1050A>G pseudoexon were evaluated using eSkip-Finder (2′OMe). High-scoring ASOs are shown in purple. The displayed threshold corresponds to the score cutoff that maximized MCC on this dataset and is shown for descriptive comparison only; primary benchmark comparisons in the main text rely on AUROC, which evaluates ranking without selecting a model-score cutoff. Because the eSkip-Finder (2′OMe) model was trained only on exon-targeting ASOs, predictions are available only for ASOs fully contained within the pseudoexon.

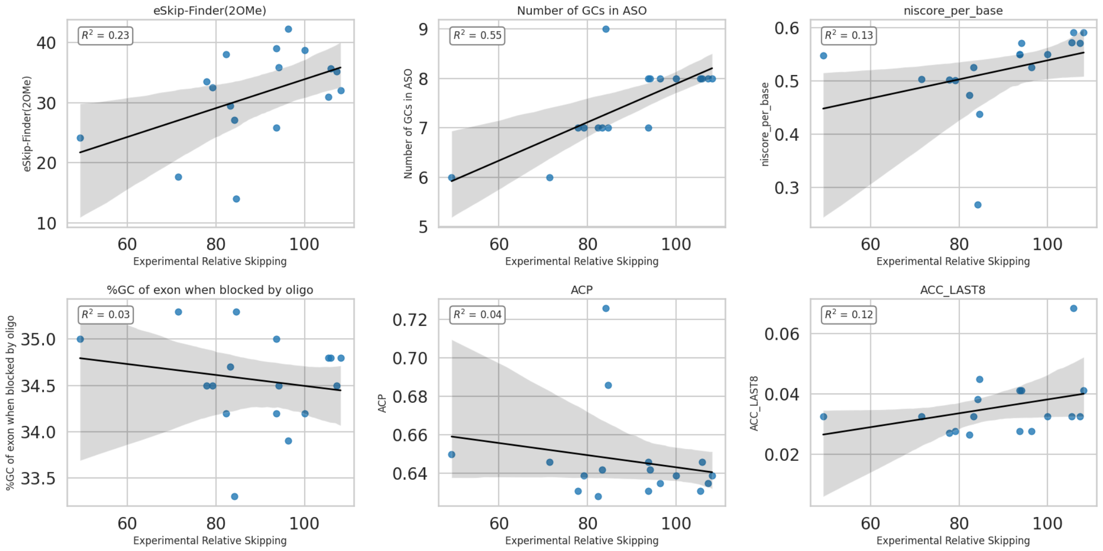


**Supplementary Figure 2 | Relationship between ATM pseudoexon activity and eSkip-Finder scores or handcrafted sequence features.**

Scatter plots show the relationship between experimental relative skipping (*x*-axis) and either the eSkip-Finder (2′OMe) score or individual handcrafted features (*y*-axis) for ASOs designed within the ATM c.5763–1050A>G pseudoexon. The plotted features are the number of GC nucleotides within the ASO, niscore_per_base, GC content of the blocked exon region, ACP (distance from the splice acceptor to the center of the target site) and ACC_LAST8 (predicted mean accessibility score for the last eight nucleotides at the 3′ end of the ASO-binding site). Black lines indicate linear regression fits and shaded regions denote 95% confidence intervals. $R^2$ values are shown in the upper-left corner of each panel.

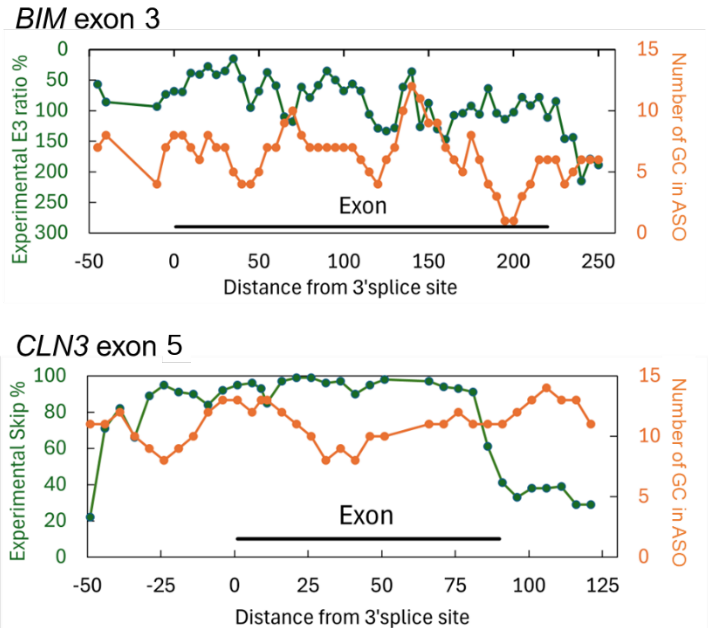


**Supplementary Figure 3 | Local GC content does not explain tiled-screen activity in BIM exon 3 and CLN3 exon 5.**

For BIM exon 3 and CLN3 exon 5, 18-nt MOE ASOs were designed in a tiled manner across the target regions, allowing a minimally biased comparison between local GC content and measured exon-skipping efficiency. The green line (left y-axis) shows the experimental E3 ratio for BIM exon 3 and exon-skipping efficiency for CLN3 exon 5, while the orange line (right *y*-axis) shows the number of GC nucleotides within each ASO. The *x*-axis indicates the distance from the 3′ splice site, defined as the distance between the most upstream nucleotide of the ASO-binding site and the 3′ splice site. Exonic regions are indicated by horizontal black bars.

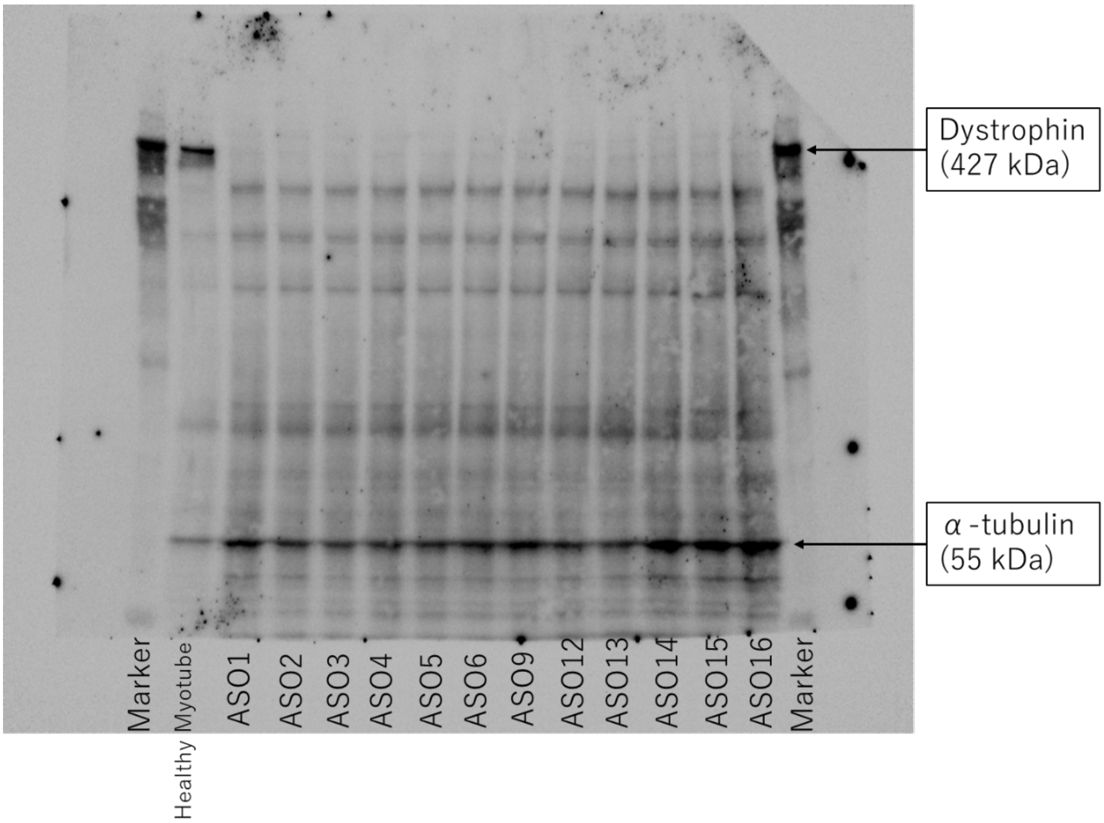


**Supplementary Figure 4 | ASOs with low exon-skipping activity do not produce detectable dystrophin restoration in *MYOD1*-converted urine-derived cells.**

Immunoblot membranes showing dystrophin expression in *MYOD1*-converted urine-derived cells from three patients with DMD carrying an exon 45 deletion after treatment at 10 μM with 12 ASOs that had each produced <20% exon skipping at 1 μM. None of these ASOs produced a detectable dystrophin signal. Molecular-weight markers are shown on both sides of the membrane. The lower band corresponds to α-tubulin as a loading control.

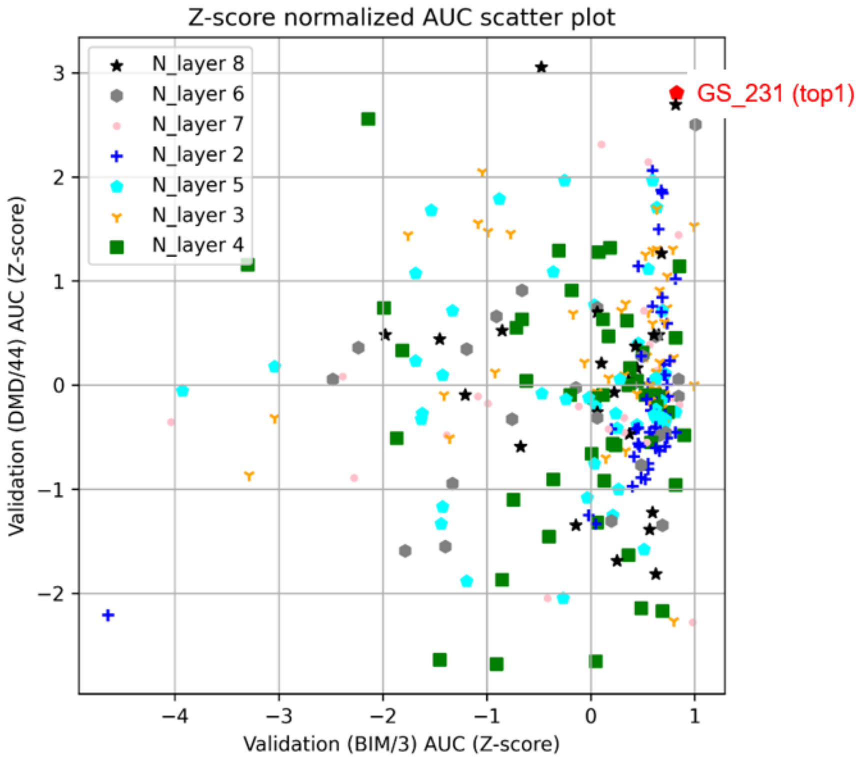


**Supplementary Figure 5 | Hyperparameter optimization for eSkip2-base.**

Z-score-normalized AUROC scatter plot comparing validation performance on BIM exon 3 (*x*-axis) and DMD exon 44 (*y*-axis). Each point represents one model from the hyperparameter search, grouped by the number of transferred network layers (2–8). The top model, GS_231, was selected a priori for the fixed benchmark and prospective analyses.

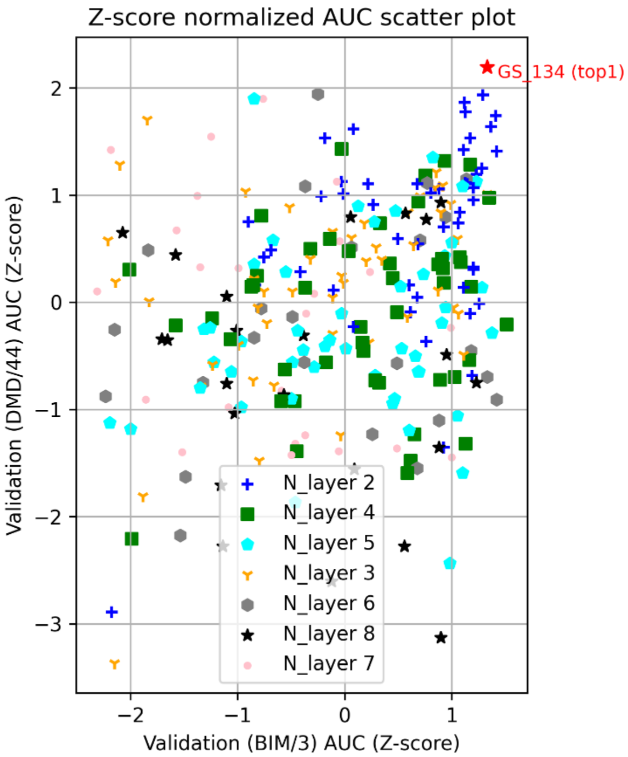


**Supplementary Figure 6 | Hyperparameter optimization for the SNV-only model (No ASO).**

Z-score-normalized AUROC scatter plot comparing validation performance on BIM exon 3 (*x*-axis) and DMD exon 44 (*y*-axis) for models trained without ASO-derived data. Each point represents one hyperparameter setting, grouped by the number of transferred layers (2–8). GS_134 was selected for subsequent analyses.

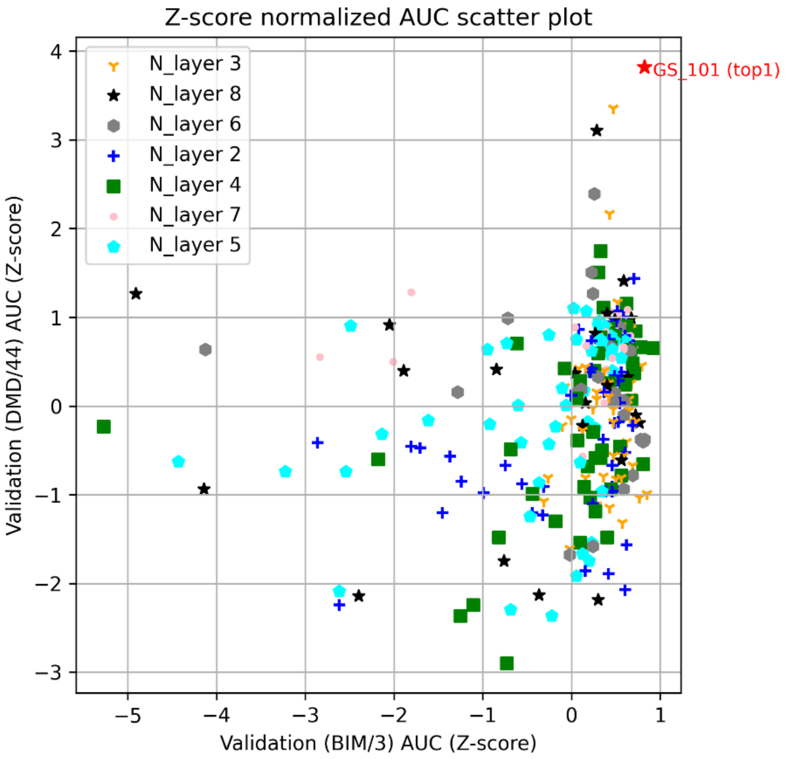


**Supplementary Figure 7 | Hyperparameter optimization for the ASO-only model (No SNV).**

Z-score-normalized AUROC scatter plot comparing validation performance on BIM exon 3 (*x*-axis) and DMD exon 44 (*y*-axis) for models trained without SNV-derived data. Each point represents one hyperparameter setting, grouped by the number of transferred layers (2–8). GS_101 was selected for subsequent analyses.

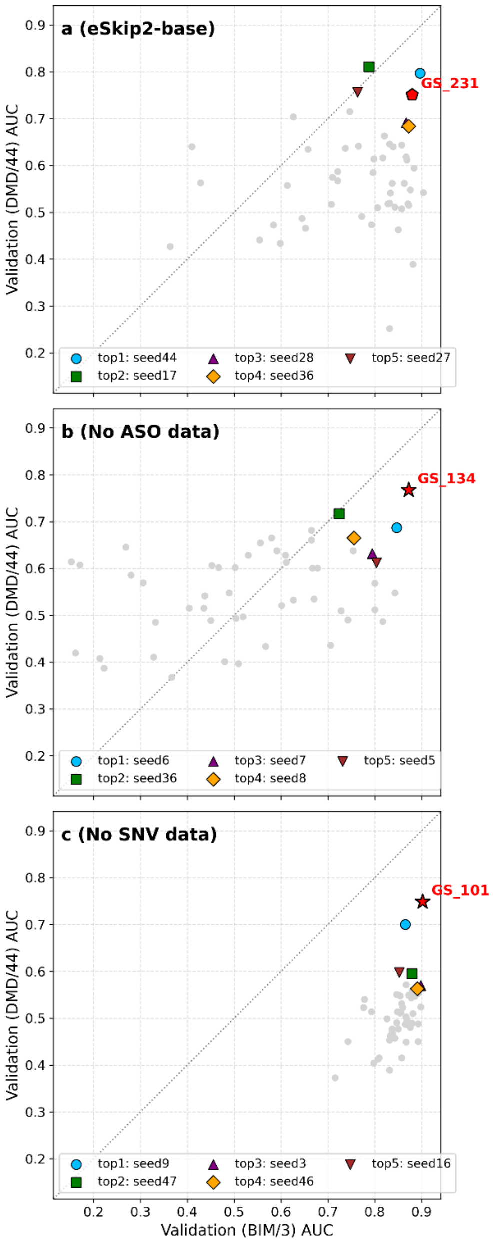


**Supplementary Figure 8 | Seed-wise validation performance distributions for eSkip2-base and single-source training variants.**

Scatter plots show validation AUROC for 50 independently trained models with different random seeds under the selected hyperparameter setting for eSkip2-base (**a**), the SNV-only model (**b**), and the ASO-only model (**c**). Each gray point represents one seed, plotted using AUROC on BIM exon 3 (*x*-axis) and DMD exon 44 (*y*-axis). Colored symbols indicate the top five seeds ranked by the validation criterion within each 50-seed run set. Red symbols denote the pre-specified runs selected during the original hyperparameter search (GS_231, GS_134, and GS_101) for reference. Dashed diagonal lines indicate equal AUROC on the two validation tasks.

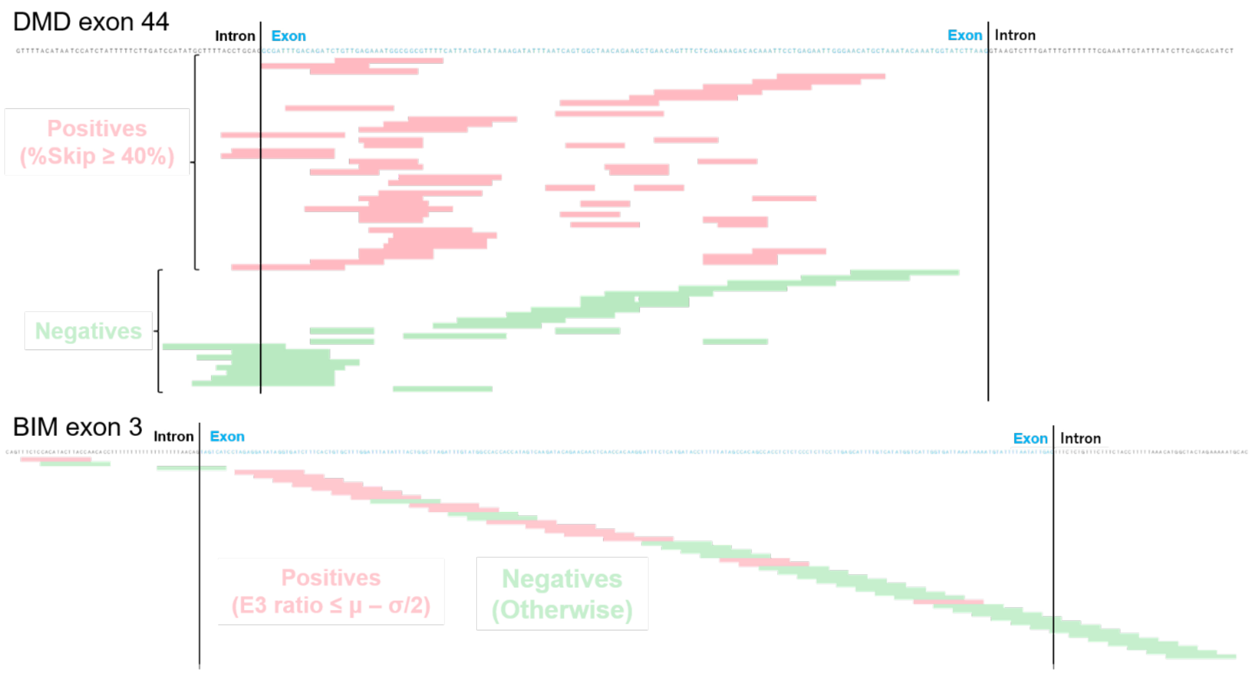


**Supplementary Figure 9 | Experimental ASO activity in the two validation datasets used for model selection.**

ASO binding sites are shown for DMD exon 44 (top) and BIM exon 3 (bottom), with exonic and flanking intronic sequences indicated in blue and black, respectively. The DMD exon 44 dataset included ASOs targeting the 3′ splice site and dual-targeting designs; ASOs with ≥40% exon skipping were labeled positive, and those with <40% skipping were labeled negative. The BIM exon 3 dataset comprised fixed-length 18-nt ASOs broadly tiled across exon-proximal sequences. Because the BIM assay measured retained-exon abundance rather than exon-skipping percentage, ASOs with E3 ratios $\leq \mu - \sigma/2$ were labeled positive, where $\mu$ and $\sigma$ denote the dataset mean and standard deviation, respectively; lower E3 ratios indicate greater exon 3 skipping. Samples near the threshold were excluded as ambiguous, and the remaining analyzed ASOs were labeled negative. Pink and green bars indicate positive and negative validation labels, respectively. The two datasets were used jointly to select models with good performance across distinct ASO architectures and assay contexts.

**Supplementary Table 1. Primary benchmark performance of eSkip2 and comparison with eSkip-Finder**

| **Region** | **Exon and intron region** | | | | | | | | **Exon region** | | |
|---|---|---|---|---|---|---|---|---|---|---|---|
| **Model** | **eSkip2** | | | | | | | | **eSkip2** | **eSkip-Finder (PMO)** | **eSkip-Finder (2′OMe)** |
| Data name (Gene/Exon) | AUROC | MCC | Accuracy | Precision | TP | TN | FP | FN | AUROC | AUROC | AUROC |
| COL7A1/73 | 0.94 | 0.86 | 0.92 | 0.85 | 17 | 19 | 3 | 0 | 0.95 | 0.51 | 0.29 |
| COL7A1/80 | 0.71 | 0.55 | 0.81 | 0.6 | 3 | 10 | 2 | 1 | Not available owing to the limited number of ASO (*n* = 1) data calculable by eSkip-Finder. | | |
| CLN3/5 | 0.85 | 0.81 | 0.91 | 0.83 | 10 | 21 | 2 | 1 | 0.63 | 0.69 | 0.34 |
| CLN3/5 mean | 0.77 | 0.65 | 0.79 | 1.0 | 16 | 11 | 0 | 7 | Not available owing to the absence of negative samples in regions calculable by eSkip-Finder. | | |
| SCN1A/ pseudoexon | 0.76 | 0.52 | 0.74 | 0.67 | 12 | 8 | 6 | 1 | 0.83 | 0.72 | 0.72 |
| ATM c.5763-1050A>G/ pseudoexon | 0.76 | 0.66 | 0.82 | 0.75 | 15 | 7 | 5 | 0 | 0.36 | 0.57 | 0.73 |
| NF1/17 | 0.76 | 0.73 | 0.85 | 1.0 | 5 | 6 | 0 | 2 | 0.76 | 0.74 | 0.41 |

AUROC is the primary benchmark metric. MCC, accuracy, precision, and confusion-matrix counts are shown only as descriptive summaries because they depend on the selected model-score cutoff. For each benchmark dataset, the cutoff was selected to maximize MCC.

**Supplementary Table 2. Training set used for the development of eSkip2**

| Gene/Exon | Positive | Negative | Reference |
|---|---|---|---|
| DMD/2 | 2 | 37 | WO2020023688A1[1] |
| DMD/45 | 60 | 94 | JP2017163994A[2]<br>JP2018183178A[3]<br>JP2020114215A[4]<br>JP2022058379A[5]<br>TW201811807A[6] |
| DMD/50 | 1 | 0 | JP2022058379A[5] |
| DMD/51 | 2 | 1 | WO2015137409A1[7] |
| DMD/53 | 34 | 93 | JP2018027083A[8]<br>JP6867636B1[9]<br>US20170369875A1[10]<br>US2014315977A1[11]<br>US2015361428A1[12]<br>US2021102205A1[13] |
| DMD/55 | 9 | 21 | JP2017163994A[2]<br>JP2020114215A[4]<br>JP2022058379A[5] |
| MSTN/2 | 141 | 73 | WO2017047741A1[14] |
| Single Nucleotide Variant (MFASS) | 1046 | 26,630 | Chong et al. 2019[15] |
| Wild type of SNV (MFASS) | 0 | 2,339 | Chong et al. 2019[15] |
| Total | 1,295 | 29,288 | |

Positive and negative labels were defined according to the data source. For ASO-derived data, records with exon-skipping efficiency ≥40% were labeled positive and those with <40% were labeled negative. For MFASS-derived data, an SNV site was labeled positive if any tested substitution at that site induced exon deletion; otherwise, it was labeled negative. The corresponding wild-type sequences were labeled negative.

**Supplementary Table 3. Validation set used for model selection**

| Gene/Exon | Label definition | Positive | Negative | Reference |
|---|---|---|---|---|
| DMD/44 | %skip ≥ 40% | 40 | 23 | JP2017163994A[2]<br>JP2018027087A[16]<br>JP2020114215A[4]<br>JP2022058379A[5]<br>US20140323544A1[17] |
| BIM/3 | E3 ratio ≤ $\mu$ - σ/2 | 18 | 30 | Liu et al. 2017[18] |

**Supplementary Table 4. Benchmark test set used for cross-gene evaluation**

| Data name (Gene/exon number) | Label definition[a] | Positive | Negative | Reference |
|---|---|---|---|---|
| COL7A1/73 | %skip ≥ 40%[b] | 17 | 22 | US20180216106A1[19] |
| COL7A1/80 | %skip ≥ $\mu$ (33%) | 4 | 12 | JP2018515122A[20] |
| CLN3/5 | %skip ≥ 95%[c] | 11 | 23 | Centa et al. 2020[21] |
| CLN3/5 mean | %skip ≥ $\mu$ (77%)[c] | 23 | 11 | Centa et al. 2020[21] |
| SCN1A/ pseudoexon 20 | %skip ≥ $\mu$ (65%) | 13 | 14 | Han et al. 2020[22] |
| ATM (c.5763-1050A>G)/ pseudoexon | Relative skip[d] ≥ $\mu$ (0.81) | 15 | 12 | Kim et al. 2023[23] |
| NF1/17 | %skip ≥ $\mu$ (20%) | 7 | 6 | Leier et al. 2022[24] |

[a] Operational thresholds were used to derive binary benchmark labels from heterogeneous experimental readouts.

[b] For COL7A1 exon 73, exon-skipping activity was originally reported as categorical ranges (0, 1–10, 11–20, …, 61–70%). The threshold (40%) corresponds to the approximate mean of these discrete categories.

[c] In the CLN3/5 experiments, several ASOs targeted intronic regions (14/34). A stringent 95% threshold was used for the primary binary benchmark to avoid labeling nearly all exon-proximal ASOs as positive; mean-based results are provided as a sensitivity analysis in Supplementary Table 1.

[d] ASO activity was reported as values normalized to the baseline ASO AT043.

**Supplementary Table 5. Experimental conditions represented across the development and benchmark datasets**

| Gene/Exon | Cell type | ASO cell entry | ASO Chemistry | ASO concentration |
|---|---|---|---|---|
| DMD/2, 44, 45, 50, 51, 53, 55 | RD cells | Electroporation or transfection | PMO<br>2′OMe | 10-12.5 μM<br>0.1 μM |
| MSTN/2 | RD cells | Transfection | 2′OMe | 0.1 μM |
| BIM/3 | K562 CML cells | Transfection | MOE | 2 μM |
| CLN3/5 | Fibroblasts derived from patients with CLN3 Batten disease | Transfection | MOE | 0.1 μM |
| COL7A1/73 | HeLa | Transfection | 2′OMe | 0.1 μM |
| COL7A1/80 | Primary human fibroblasts | Transfection | 2′OMe | 0.1 μM |
| SCN1A/ pseudoexon | ReNcells VM | ASO gymnotic (free) uptake | MOE | 20 μM |
| ATM (c.5763-1050A>G)/ pseudoexon | Fibroblasts derived from patients with AT (c.5763-1050A>G) | Transfection | MOE | 0.2 μM |
| NF1/17 | HEK293 | Transfection | PMO | 2 μM |

**Supplementary Table 6. Prospectively tested DMD exon 46 candidates: experimental skipping efficiencies and eSkip2 scores**

| | Skipping efficiency % | s.d. % | eSkip2 score | | | Score rank (Max) |
|---|---|---|---|---|---|---|
| | | | Binding region (12x2) | Binding region (13+11) | Max | |
| Non-treated | 3 | 2 | | | | |
| #1 | 0 | 0 | 0.99879 | | 0.99879 | 5 |
| #2 | 0 | 0 | 0.99880 | 0.99887 | 0.99887 | 4 |
| #3 | 11 | 16 | 0.99837 | | 0.99837 | 9 |
| #4 | 0 | 0 | 0.99877 | | 0.99877 | 6 |
| #5 | 0 | 0 | 0.99870 | | 0.99870 | 7 |
| #6 | 0 | 0 | 0.99893 | | 0.99893 | 2 |
| #7 | 30 | 12 | 0.99895 | 0.99922 | 0.99922 | 1 |
| #8 | 50 | 3 | 0.99892 | | 0.99892 | 3 |
| #9 | 0 | 0 | 0.85401 | | 0.85401 | 12 |
| #10 | 19 | 6 | 0.93206 | | 0.93206 | 11 |
| #11 | 21 | 3 | 0.65679 | | 0.65679 | 15 |
| #12 | 4 | 6 | 0.67918 | | 0.67918 | 14 |
| #13 | 0 | 0 | 0.99868 | | 0.99868 | 8 |
| #14 | 8 | 7 | 0.99538 | | 0.99538 | 10 |
| #15 | 14 | 15 | 0.77976 | | 0.77976 | 13 |
| #16 | 7 | 8 | 0.43375 | | 0.43375 | 16 |
| Positive control (30-mer) | 89 | 14 | | | | |

**Supplementary Table 7. Hyperparameter search space for eSkip2-base**

| Setting of Fine-Tuning for Exon Skipping | Range |
|---|---|
| Pretrained HyenaDNA model | Pretrained model using a 160k sequence window. "hyenadna-medium-160k-seqlen" |
| Number of layers | 2, 3, 4, 5, 6, 7, 8 |
| Batch size | 256, 512 |
| Width | 256 |
| Sequence length | 500 |
| Embed dropout | 0, 0.1, 0.2, 0.3 |
| Weight decay (model) | 0.1 |
| Weight decay (hyena layer) | 0 |
| Resid dropout | 0 |
| Reverse complement augmentation | False |
| Learning rate | 3e-5, 1e-4 |
| Learning rate schedule | Warm Up and Cosine Decay |
| Epoch | 25, 50, 100 |
| Label smoothing | 0.01 |
| Floating-point formats | TensorFloat-32 |
| Number of parameters for the sequence decoder | 514 |

**Supplementary Table 8. Selected hyperparameters for eSkip2-base**

| Setting of fine-tuning for exon skipping | GS_231 |
|---|---|
| Number of layers | 5 |
| Batch size | 256 |
| Embed dropout | 0.3 |
| Learning rate | 3e-5 |
| Epoch | 100 |

**Supplementary Table 9. Exon sets used for gene-adaptive refinement**

| Data name (Gene/Exon) | Exon sets used for gene-adaptive refinement | NCBI RefSeq |
|---|---|---|
| COL7A1/73 | Exon 2-117 | NG_007065.1 |
| COL7A1/80 | | |
| CLN3/5 | Exon 2-14 | NG_008654.2 |
| CLN3/5 mean | | |
| SCN1A/ pseudoexon | Exon 2-15, 17-25, pseudoexon<br>Exon 16 exceeds the model's limit. | NG_011906.1 |
| ATM (c.5763-1050A>G)/ pseudoexon | Exon 2-62, pseudoexon[23], WT of pseudoexon[23] | NG_009830.1 |
| NF1/17 | Exon 2-20, 22-36, 38-57<br>Exons 21 and 37 exceed the model's limit. | NG_009018.1 |
| DMD/46 | Exon 2-78 | NG_012232.1 |

**Supplementary Table 10. Hyperparameters for gene-adaptive refinement of eSkip2**

| Setting of gene-adaptive refinement | Value |
|---|---|
| Pretrained model | eSkip2-base |
| Number of layers | 5 |
| Batch size | 256 |
| Embed dropout | 0 |
| Resid dropout | 0 |
| Learning rate | 1e-6 |
| Learning rate schedule | Warm Up and Cosine Decay |
| Epoch | 200 |
| Adopted model | Epoch with lowest validation loss |

**Supplementary Table 11. Selected hyperparameters for the SNV-only model (No ASO)**

| Setting of fine-tuning for exon skipping | GS_134 |
|---|---|
| Number of layers | 8 |
| Batch size | 256 |
| Embed dropout | 0.2 |
| Learning rate | 1e-4 |
| Epoch | 100 |

**Supplementary Table 12. Selected hyperparameters for the ASO-only model (No SNV)**

| Setting of fine-tuning for exon skipping | GS_101 |
|---|---|
| Number of layers | 8 |
| Batch size | 256 |
| Embed dropout | 0.2 |
| Learning rate | 3e-5 |
| Epoch | 25 |
